# The dynamic gene regulatory landscape of the mouse brain in response to sleep deprivation

**DOI:** 10.1101/2024.07.04.602083

**Authors:** Akanksha Bafna, Quang Dang, Lewis Taylor, Steven Walsh, David Sims, Charlotte George, Aarti Jagannath

**Affiliations:** Sleep and Circadian Neuroscience Institute (SCNi), Nuffield Department of Clinical Neurosciences, New Biochemistry Building, University of Oxford, South Parks Road, Oxford, OX1 3QU, U.K.; Department of Physiology, Anatomy and Genetics, University of Oxford, Mansfield Road, Oxford, OX1 3PT, U.K.; Exscientia, The Schrödinger Building Oxford Science Park, Oxford, OX4 4GE; Evonetix, 9a Coldham’s Business Park, Norman Way, Cambridge, CB1 3LH; Centre for Computational Biology, MRC Weatherall Institute of Molecular Medicine, University of Oxford, John Radcliffe Hospital, Headington, Oxford, OX3 9DS

**Author notes:** Lead contact AJ.

## Abstract

Sleep deprivation (SD) negatively impacts nearly all brain functions including cognition, memory consolidation and metabolism. However, the gene regulatory networks that underlie these biological effects are not well understood. In order to identify these networks, we conducted a multiomic analysis to analyse how gene expression, chromatin accessibility, enhancer activity and DNA methylation change with acute SD in mice. By studying three brain regions involved in different aspects of sleep - the cortex (CTX), dentate gyrus (DG), and suprachiasmatic nucleus (SCN) - we found that the effects of SD on the multiome varied widely, impacting physiological processes specific to each area, from spine formation in the dentate gyrus to neuropeptide release in the SCN. Our integrated analysis showed that distinct brain region-specific networks of regulatory factors dynamically alter the epigenomic landscape in response to SD to orchestrate transcriptional responses. These findings provide new understanding of the regulatory grammar encoded within the genome, which enables a general physiological signal like SD to produce unique effects in a tissue-specific context.

## INTRODUCTION

Sleep is evolutionarily conserved, complex, and multi-faceted process that regulates the physiology of the brain and body. It is clear that the loss of sleep leads to widespread deleterious effects on cognition, learning and memory, attention and metabolism. (Hudson et al., 2020; McHill & Wright, 2017; Raven et al., 2018). Despite decades of research, the molecular determinants of sleep, and those that underlie the deleterious effects of sleep loss remain mysterious.

Furthermore, the mammalian brain shows many distinct anatomical substructures that differ markedly in their molecular composition, physiology and function in sleep. In illustration, the regulation of sleep/wake behaviour is governed by two key processes that are localised to different parts of the brain. The first is the homeostatic sleep drive or the need for sleep, where CTX and hippocampus serve as a record keepers of sleep loss (Deboer, 2018; Hastings et al., 2019; Thomas et al., 2020). The second is circadian drive which regulates the timing of sleep relative to the light/dark cycle and is localised to the hypothalamic SCN (Chiu & Prober, 2013; Edgar et al., 1993; Maywood et al., 2021; Shafer & Keene, 2021). Acute sleep loss or SD affects light sensitivity as well as circadian timing within the SCN (Jagannath et al., 2021). In contrast, sleep loss impacts memory consolidation in the hippocampus, and functional connectivity, working memory and cognition in the cortex (Hong et al., 2023; Simon et al., 2022). Such differences warrant a molecular investigation into how these brain regions respond differently to sleep loss.

Acute sleep deprivation (SD) has been the preferred method to gain molecular insights on sleep regulation. Several studies have characterised the whole brain transcriptomic, proteomic and metabolomic changes post SD in mice (Cirelli, 2006; Cirelli et al., 2004; Cirelli & Tononi, 2000; Diessler et al., 2018; Gaine et al., 2021; Mackiewicz et al., 2007; Vecsey et al., 2012). However, brain region specific differences, and the underlying molecular gene regulatory landscape are little understood. Genomic architecture is highly dynamic and plays an active role in regulating tissue/condition-specific gene expression (Fullard et al., 2018; Gray et al., 2015; Misteli, 2020; Robson et al., 2019). From DNA methylation to binding of transcription factors, multiple gene regulatory networks converge to de-encrypt genomic grammar to alter gene expression in a highly targeted manner. Therefore, a key question that remains to be answered is how the genomic architecture changes in different brain regions to achieve the response to sleep loss that is appropriate for its particular function.

To study the effect of acute sleep deprivation on the gene regulatory landscape, we conducted the simultaneous profiling of total RNA including coding and non-coding transcripts, chromatin accessibility and DNA methylation in three distinct mammalian brain regions; somatosensory cortex (CTX), dentate gyrus of the hippocampus (DG) and the suprachiasmatic nucleus (SCN) of mice. From the integrated analysis of all the readouts, we identified genome-wide but brain-region specific networks of regulatory factors that orchestrate the hierarchical and coordinated transcriptional response to sleep loss.

## RESULTS

### Sleep deprivation changes the transcriptome of the brain in a region-specific **manner**

To study how acute sleep deprivation differentially impacts the global transcriptome of distinct regions of mammalian brain, we sleep deprived mice by exposure to novel objects for 6h (ZT 0-6) and collected tissue punches from three regions of the brain with distinct roles in the regulation of sleep and responses to sleep loss. These were CTX where sleep loss impairs functional connectivity, the DG of the hippocampus where sleep regulates memory consolidation, and the SCN as the hypothalamic centre which controls the circadian timing of sleep, where sleep loss regulates the sensitivity of the circadian clock to environmental input (Jagannath et al., 2017; Jagannath et al., 2021). We compared gene expression between the brain regions under sham conditions, and in response to SD. The first comparison (between regions under sham conditions) highlighted genes that confer tissue specificity, such as the transcription factor *Zfhx3* (Zinc Finger Homeobox 3) in the SCN (Parsons et al., 2015; Wilcox et al., 2021), *Camk2n1* (Calcium/Calmodulin Dependent Protein Kinase II Inhibitor 1) in the CTX (Ling et al., 2011) and *C1ql2* (Complement C1q like 2) which regulates the number of excitatory synapses in the DG (Fig. 1a,b,c). Principal component analysis (PCA) showed that the transcriptome clustered clearly by brain region (Supplementary Fig. 1). Interestingly, the majority of the detected transcriptome was region-specific, for example 2,289 genes were robustly differentially expressed between CTX and SCN (p adjusted < 0.01, log_2_FC (fold change) ≥ 1, Supplementary Table 1), confirming substantial heterogeneity in gene expression in different parts of the brain. These tissue-specific genes were associated with biological processes known to be specific for each region. For example synaptic plasticity, learning and memory were enriched in CTX and DG while neuropeptide signalling (Ono et al., 2021), cilium movement (Tu et al., 2023) were noted in the SCN (Fig. 1e). We then segmented the differential genes in a brain region-specific manner to understand which genes were globally dysregulated following SD, and which ones showed brain-region specificity. The tissue that showed the greatest transcriptional change following SD was CTX (n = 2285), followed by DG (n = 1911) and the SCN (n = 896) at p adjusted < 0.1 (Fig. 2a-c). Of these however, only 209 genes changed across all three tissues reflecting the underlying heterogeneity in baseline gene expression. Amongst these 209 genes were those that have been described previously to be responsive to SD. For example, we found a reduction in the expression levels of *Cirbp* (cold-inducible RNA-binding protein) after SD (Hoekstra et al., 2019) uniformly across all the three tested tissues (Fig. 2f). In contrast, *Homer1* (homer scaffolding protein 1), whose mRNA levels have been reported by multiple studies to correlate with wakefulness in the brain (Diering et al., 2017; Zhu et al., 2020) was upregulated following SD in DG and CTX, but not in the SCN (Fig. 2d, f), suggesting there are brain region-specific regulatory elements at play.

**Figure 1.**
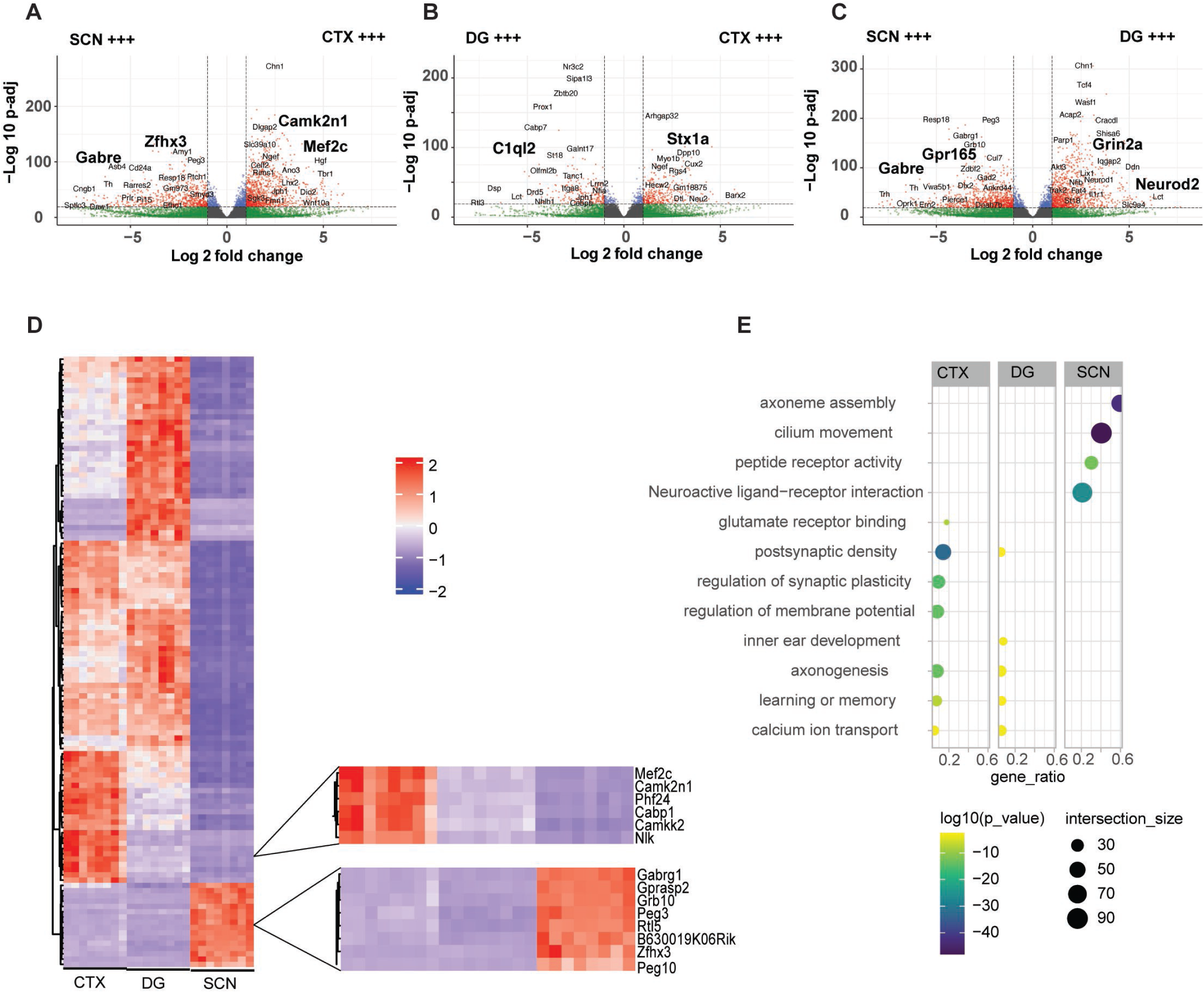
Region-specific gene expression profiles of the mouse brain. *(A,B,C)* Volcano plot log_2_ fold change and logarithmic P-value (-log_10_P) for differential gene expression between somatosensory cortex (CTX), dentate gyrus (DG) and suprachiasmatic nucleus (SCN). Red indicates those transcripts that show a log_2_ fold change > 0.5 and also a p-value < 10^-5^. *(D)* Clustered heat map view of differential genes as observed between CTX, DG and the SCN, selected clusters enlarged for genes of interest that characterise each brain region. *(E)* gProfiler plot showing enriched functional categories (y-axis) for the differential genes as per gene ontology (biological process, molecular functions and cellular component) and KEGG pathway analysis for cortex, DG and the SCN.

**Figure 2.**
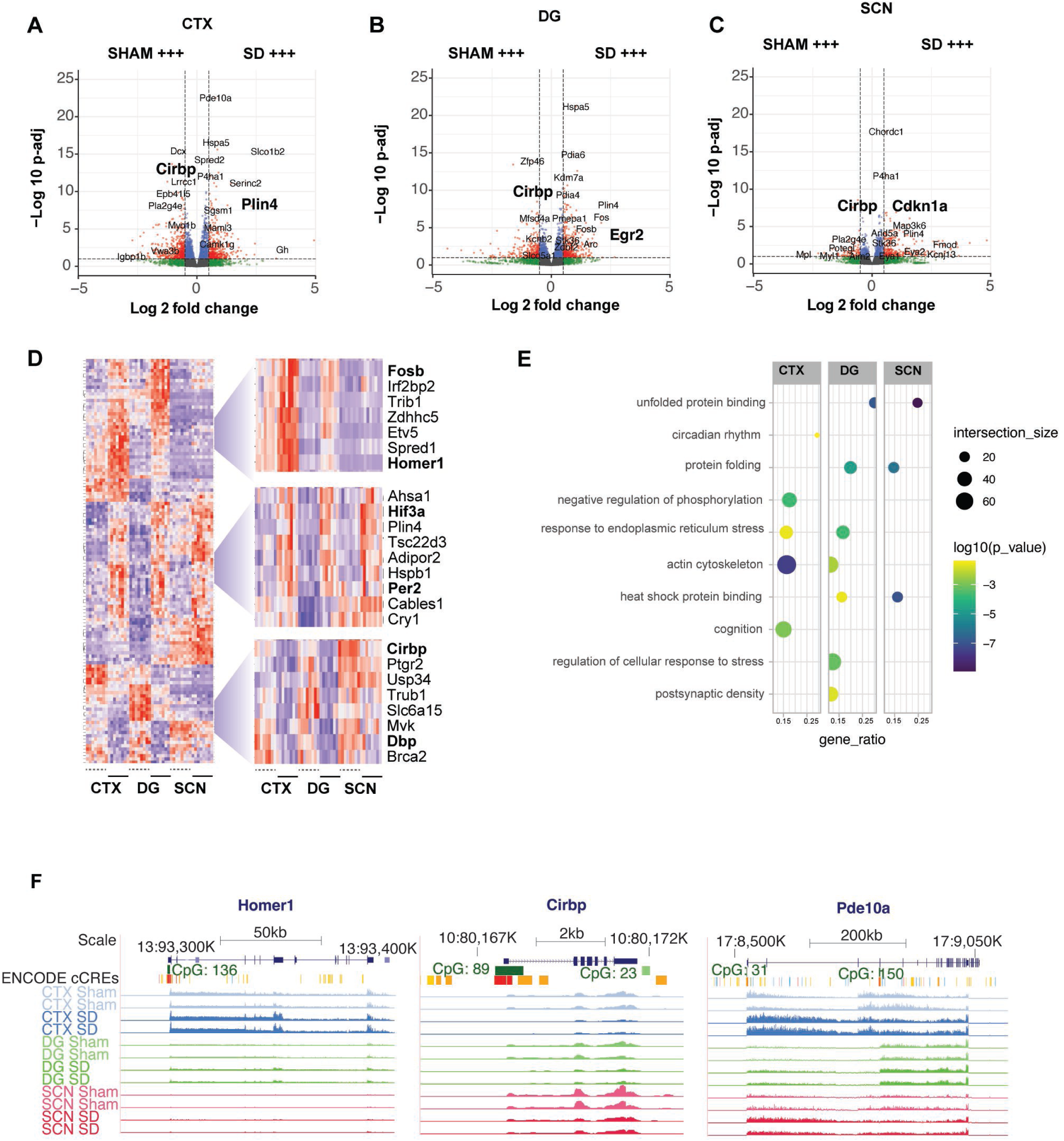
Effect of Sleep deprivation on gene expression in distinct brain regions. *(A)* Volcano plot showing Log_2_ fold change and logarithmic P-value (-log_10_P) for differential gene expression between SHAM and sleep-deprived (SD) mice in Cortex (CTX), *(B)* Dentate gyrus (DG) and *(C)* Suprachiasmatic nucleus (SCN). *(D)* Clustered Heat map view of differential genes (SHAM vs SD) in CTX, DG and SCN, clusters with genes of interest enlarged in the right. The top cluster illustrates genes that are upregulated significantly in the CTX, the second shows those that are upregulated in all three and the third genes that are downregulated. *(E)* gProfiler plot showing enriched functional categories (y-axis) for the differential genes between sham and SD as per gene ontology (biological process, molecular functions and cellular component) and KEGG pathway analysis for cortex, DG and the SCN *(F)* UCSC Genome Browser tracks of normalized RNA for Sham and SD conditions at *Homer1, Cirbp* and *Pde10a* gene loci as observed in CTX (blue), DG (green) and the SCN (pink). The chromosome location and scale (mm10 genome) are indicated at the top along with ENCODE tracks for cCRE elements and CpG methylation.

We conducted functional annotation of the genes that changed after SD for each brain region (CTX, DG, SCN) using gene ontology and pathway analysis on GProfiler2 (Kolberg et al., 2020). Genes involved in phosphorylation, regulation of metabolic processes, catalytic activity etc. were seen to be affected uniformly across all the three tested tissues indicating a generic effect in response to acute sleep loss. However, specific terms such as post-synaptic density and cognition were seen to be affected in DG and CTX, respectively, but not in the SCN (Fig. 2e). Moreover, circadian rhythm genes (such as *Per2*, *Cry1*) were deregulated in CTX and DG with larger fold-changes than in the SCN (central pacemaker), supporting the idea that local clocks in the brain are modified by sleep-wake rhythms (Abe et al., 2001; Dudley et al., 2003), and do not engage in self-sustained circadian rhythms like the SCN. Furthermore, previous studies have shown that the SCN is relatively shielded from physiological cues such that photic input remains the strongest signal (Dudley et al., 2003; Le Minh et al., 2001; Wakamatsu et al., 2001). For example, glucocorticoids and sleep deprivation both have a smaller effect on clock gene expression within the SCN (Hor et al., 2019), and this is reflected in our findings (Fig. 2).

We then considered those genes that were differentially expressed in one region only (Methods). This analysis found approximately 477 genes that were differential in the CTX alone, versus 146 that were significant in the SCN. The specific changes again reflected the underlying biological function of the respective brain regions. In illustration, we found that neuropeptides such as *Vip* (Vasoactive Intestinal Peptide) and *Npy* (Neuropeptide Y) which are known to be involved in circadian timekeeping are differentially regulated in response to SD in the SCN only (Fukuhara et al., 2001; Ono et al., 2021). In contrast, those involved in learning and memory such as *Atxn1, Slc1a1* (Afshari et al., 2017; Handler et al., 2023) are disrupted in CTX. This region-specific change in gene expression suggests a coordinated gene regulatory landscape specific to each brain region that underlies a distinct response to SD.

### Sleep deprivation changes the accessible chromatin landscape in a brain **region-specific manner.**

To identify the upstream drivers of the region-specific transcriptional response to SD, we used Assay for Transposase-Accessible Chromatin sequencing (ATAC-Seq) (Bao et al., 2015; Buenrostro et al., 2015) and profiled open chromatin regions in CTX, DG and SCN collected from sham control and SD mice under identical conditions to the RNA-Seq. A total of 150,234 open chromatin regions (OCRs) were detected across the CTX, DG and the SCN brain regions (Methods). We analysed region-specific OCRs by conducting differential peak analysis (Diffbind, FDR ≤ 0.05, Supplementary Table 2) between the tissues. In the comparison between CTX and SCN, ca. 42% of the differential OCRs mapped to distal intergenic regions, with < 20% mapping within 3kb of the promoter (Fig. 3a). Each OCR peak was then assigned to the nearest neighbouring target gene (TSS; transcription start site). As expected, genes that were transcribed highly in specific brain regions, such as *Camk2a* (cortex, DG) (Frankland et al., 2001) and *Six6* (SCN) (Clark et al., 2013) were associated with differential tissue-specific OCRs (Fig. 3b) at or near the promoter. However, given there were multiple proximal and distal OCRs within and around each gene, we noted that altered transcription was not always associated with changed accessibility at the promoter. In illustration, in the case of the GABA type A receptor subunit epsilon, *Gabre,* which is SCN-specific in expression, we observed no differential peaks at the promoter. This prompted us to assess all OCRs within ± one million bases of TSS corresponding to tissue-specific genes (from RNA sequencing). A high correlation was noted between tissue-specific gene expression and closest chromatin accessibility in certain cases, in other cases the strongest correlation was observed with a distal element. As exemplified by the *Gabre* gene, the strongest correlated OCR is about 800,000 bp upstream (correlation coefficient = 0.99, Fig. 3c, Supplementary Table 3), which is accessible in the SCN alone. On further inspection for TF binding sites using JASPAR (Rauluseviciute et al., 2024), the distal element showed enrichment for RFX (Regulatory factor X4) factor binding which is postulated to be important for light signalling in the SCN (Araki et al., 2004). Thus, chromatin accessibility does define gene regulation but the closest accessible region and changes at the promoter may not always be responsible for transcript regulation. This is further supported by examining the strength of association between chromatin accessibility and closest gene expression levels across the distinct brain regions (Supplementary Fig. 2). As expected, we noted almost equal distribution of number of OCRs demonstrating high (Pearson > 0.5) and low (Pearson < 0.5) correlation with tissue-specific gene expression.

**Figure 3.**
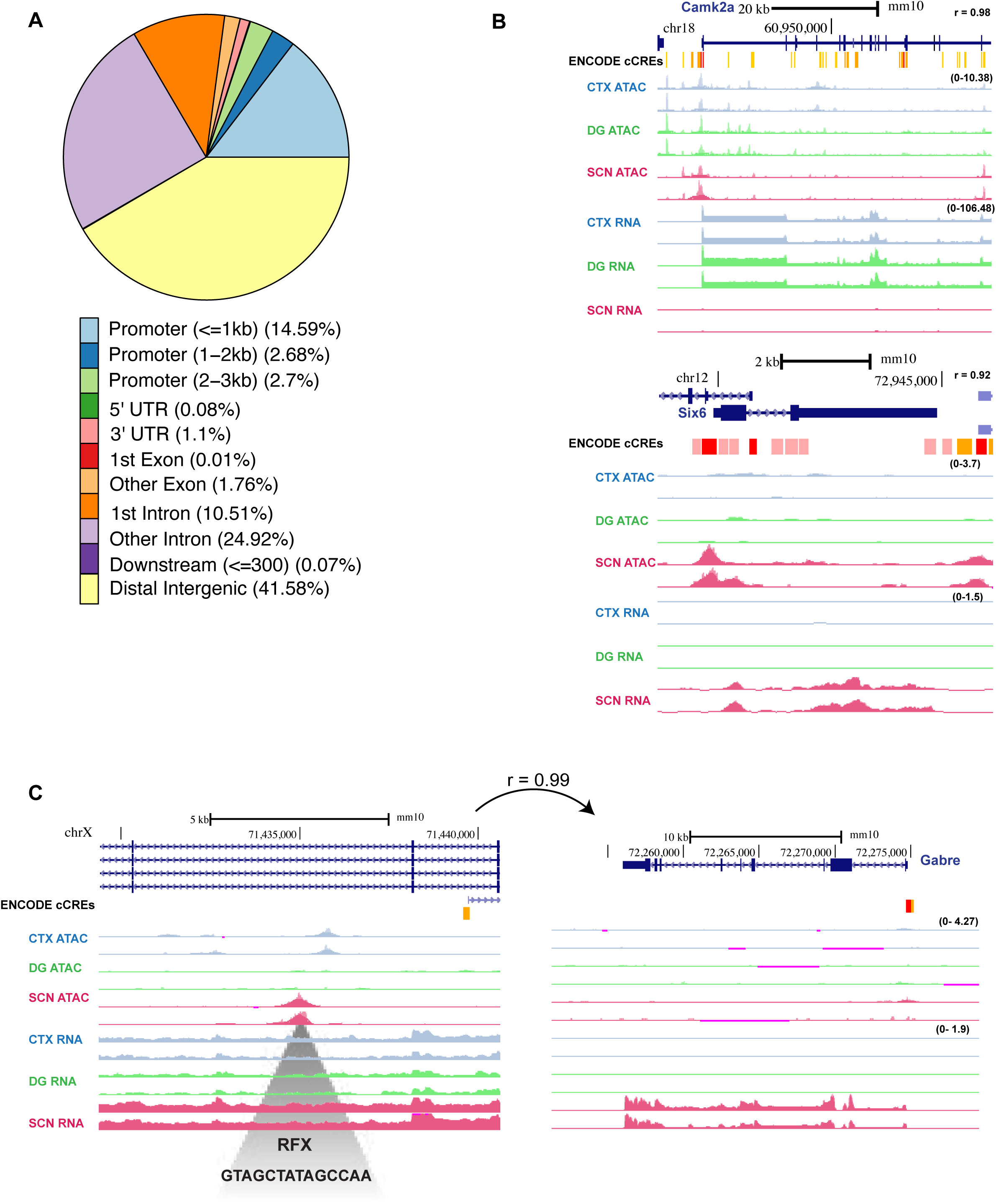
Region-specific accessible chromatin regions in the mouse brain. *(A)* Genomic feature distribution of differential OCRs between CTX and the SCN as computed by ChIPSeeker. *(B)* UCSC Genome Browser tracks for normalized RNA and accessible chromatin (ATAC-seq coverage) as indicated at *Camk2a, Six6* and *(C) Gabre* gene locus in CTX (blue), DG (green) and SCN (pink). The chromosome location and scale (mm10 genome) are indicated at the top along with ENCODE tracks for cCRE elements. Note no accessible peaks at the *Gabre* promoter, instead an upstream distal OCR that correlated (r = 0.99) with *Gabre* expression is illustrated. The grey shaded region represents RFX binding site using JASPAR.

We then interrogated each brain region for differential chromatin accessibility driven by sleep-deprivation, by comparing sham and SD peaks. Genes showing a global (in all tested regions) or region-specific response to SD in their expression levels were compiled, and their relationship was assessed with mapped OCRs (Supplementary Table 4). Reflecting the findings with tissue-specific OCRs, SD responsive OCRs also showed different patterns of correlations with their proximal genes. Together, these results suggest that a one-to-one correlation between OCRs and gene expression does not exist and should be considered on a case-by-case basis. For example, hypoxia inducible factor 3 alpha subunit (*Hif3a*) showed both increased promoter accessibility and gene expression following sleep deprivation, indicating a promoter-driven ubiquitous regulation in response to SD (Fig. 4a). However, for the well-characterised SD responsive gene *Cirbp* (cold inducible RNA binding protein), the promoter OCR did not correlate with the decreased expression levels after SD (Fig. 4b), but rather showed increased accessibility (particularly in the DG). This opening of chromatin upstream of *Cirbp* gene in response to SD might result in transcriptional repression through the binding of a repressor. Similarly, *Homer1* (homer scaffolding protein 1) showed a clear increase in transcript within the CTX and DG (not in the SCN) but the peak of accessibility at the promoter remained open under all conditions. These findings suggested the region-specific response to SD was not mediated at the promoter (Fig. 4c), but from a distal element. Indeed, we found multiple OCRs that correlated with the observed transcript levels within a million bases up/downstream of the transcriptional start site (TSS) (Supplementary Table 4). One such OCR with a high degree of correlation with transcript expression (correlation coefficient = 0.91) was found 600,000bp upstream, near the promoter of the gene *Bhmt*. Analysis of this distal element for TF binding sites using JASPAR (Rauluseviciute et al., 2024) identified several putative TF binding sites, of which KLF4 (Krüppel-Like Factor) had the highest score (14.5 by JASPAR), and KLF is known to be involved in cortical and hippocampus development (Li et al., 2021). Taken together, these data support a role for enhancers and repressors acting at distal regulatory elements, as much as transcriptional activators in shaping dynamic gene regulation in response to SD. JASPAR analysis of multiple region-specific OCRs indicated certain motifs that might be able to characterise each brain region or its response to SD, such as AP-1 factors, which were over-represented across all three brain regions. Therefore, in order to comprehensively capture the regulatory elements that could mediate the response to sleep-deprivation in a tissue specific manner, we decided to investigate the chromatin accessible regions for transcription factor binding and enhancer activity.

**Figure 4.**
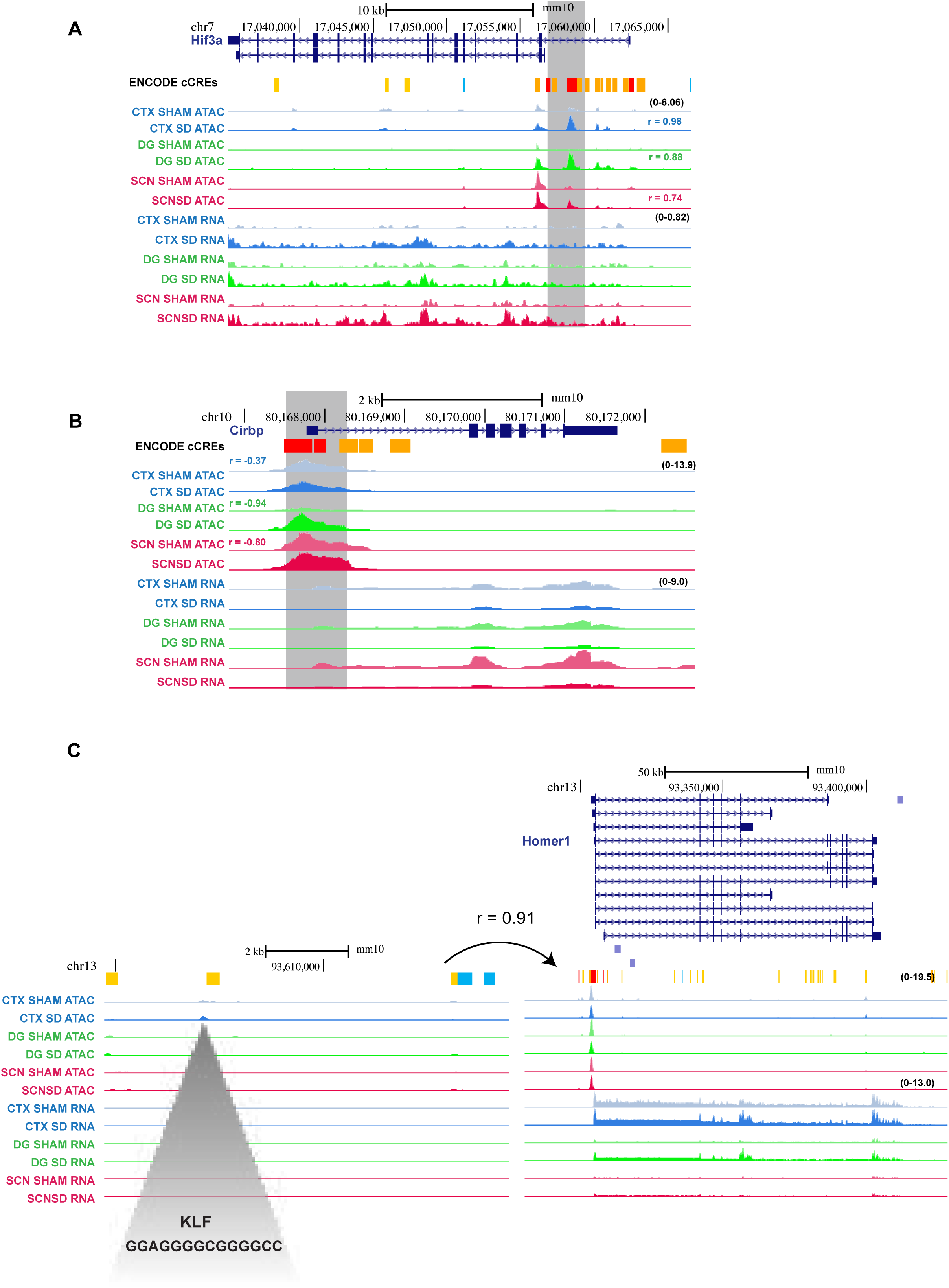
Region-specific effect on chromatin accessibility after acute SD. *(A)* UCSC Genome Browser tracks for normalized RNA and accessible chromatin (ATAC-seq coverage) for SHAM and SD conditions (values indicated on top right) for CTX (blue), DG (green) and the SCN (pink) around the *Hif3a, (B) Cirbp and (C) Homer1* gene locus illustrating distal gene regulatory element correlating with the transcriptomic response to SD in the CTX alone. The grey marked region represents KLF-4 binding site using JASPAR. The chromosome location and scale (mm10 genome) are indicated at the top along with ENCODE tracks for cCRE elements. Highlighted sites (grey) represent the SD correlated peaks with Pearson’s correlation for each tested region indicated at the top of the ATAC tracks.

### Identification of genome-wide enhanced transcription factor binding that underlie region-specific – and SD responsive-changes in chromatin accessibility

We hypothesised that the analysis of OCRs in a genome-wide fashion for overrepresented or depleted TF binding motifs could identify the key regulatory elements that shaped the transcriptome in a region and condition-specific manner. We conducted systematic transcription factor footprinting analysis as previously described (Bentsen et al., 2020) to identify which motifs were over- or underrepresented in the OCRs in the region-specific and SD-specific responses. We saw that binding sites of TFs that characterised specific tissues - RFX and MEF2C (Myocyte enhancer factor 2c) - were highly abundant in open chromatin regions found in the SCN (Araki et al., 2004) and CTX (Harrington et al., 2016), respectively (Fig. 5). This is in agreement with the finding that tissue-specific TFs determine tissue-specific gene expression (Lu et al., 2022). In particular, RFX4 has been implicated in the transcriptional response to light in the SCN, while MEF2C plays an important role shaping cortical excitatory and inhibitory synaptic connections. The steroid sex hormone receptors - Androgen receptor AR AR and Oestrogen receptors ESR1/2 - are abundantly expressed in the SCN both through development and adulthood, and this was reflected in our data as increased binding for these factors was noted in the SCN. However, it was interesting that binding sites for other steroid hormones such as NR3C1/2 were also overrepresented in the SCN, when the SCN has long been considered shielded from the effects of glucocorticoid signalling (Lehmann et al., 2023).

**Figure 5.**
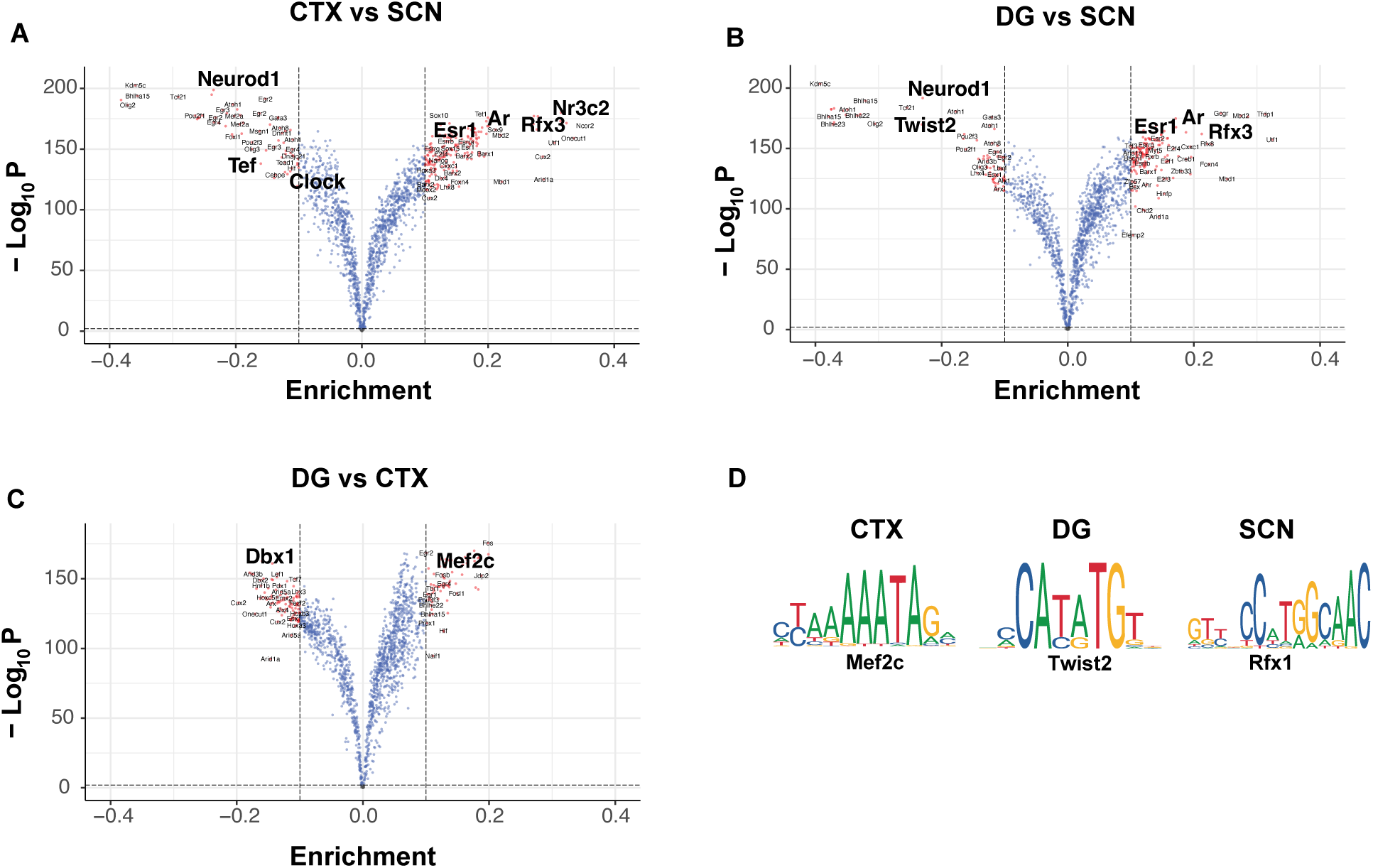
Enrichment of transcription factor binding sites in accessible chromatin regions. *(A, B, C)* Volcano plot showing enrichment and logarithmic P-value (log_10_P) for enrichment of TFs in open chromatin regions (OCRs) between distinct brain regions as computed by TOBIAS. *(D).* Associated DNA motif for the enriched region-specific TFs.

We then determined which TF binding motifs characterised the response to SD. Reflecting the transcriptomic and chromatin accessibility differences, there were fewer binding sites that characterised SD responses relative to those that specified the difference between brain regions (Fig. 6). We observed a global enrichment of immediate-early activators such as AP-1 family transcription factors (FOS, JUN) following SD across all three brain regions. However, the binding of tissue-specific transcription factors, such as MEF2C in the CTX, AR in the SCN, and TWIST2 in the DG (Fig. 6a-c) all reduced after SD. The observations at the level of TF binding are consistent with the transcriptome-level changes observed in the brain regions, where 1) the factors that bound in a region-specific manner were also expressed in the same region (Fig. 6d) 2) the upregulation of immediate-early genes AP-1 members following SD as previously described (Terao et al., 2006; Tononi & Cirelli, 2006), is seen consistently across different brain regions in our data (Supplementary Table 1, Fig. 6d).

**Figure 6.**
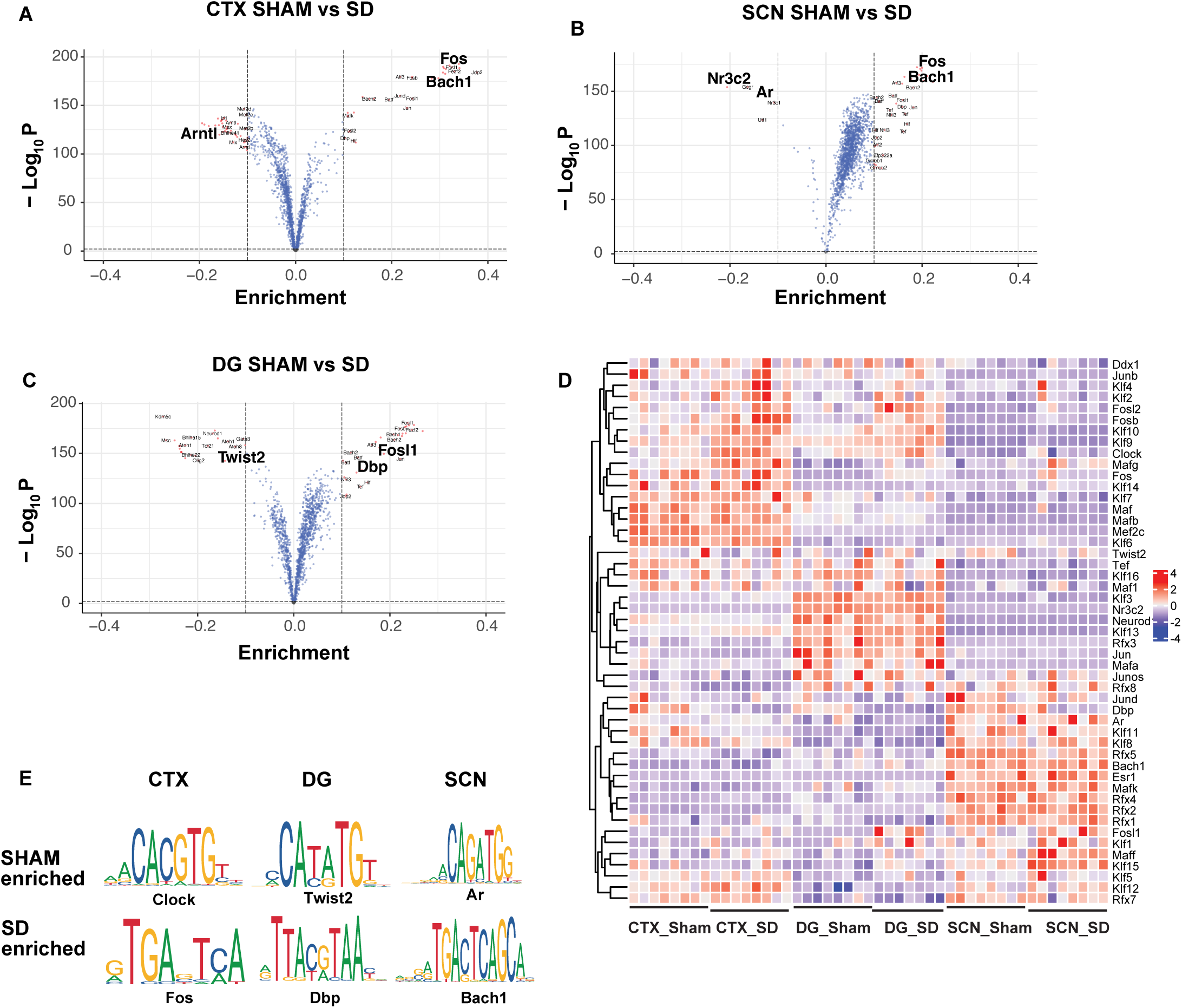
Enrichment of transcription factor binding sites under sleep-deprivation. *(A)* Volcano plot showing enrichment and logarithmic P-value (log_10_P) for enrichment of TFs in open chromatin regions (OCRs) between sham-control and sleep-deprivation (SD) mice in cortex, *(B)* Dentate gyrus (DG) and *(C)* SCN *(D)* Heatmap illustrating the normalised RNA levels of the differential TFs in Sham and SD conditions for the studied tissues. *(E).* Associated DNA motif for the enriched tissue-specific TFs in sham and SD conditions.

We then examined the co-occurrence of different TF binding sites within the same OCRs and found co-operative regulation in a region-and-condition dependent manner (Supplementary Fig. 3) for distinct families of TFs. Under SD conditions, co-occurrence of transcription factor binding sites were enriched for the FOS, JUN family in all the three tissues, and they frequently occurred with the chromatin remodelling factor ARID3A (AT-rich interaction domain 3A) to regulate target gene expression (Lin et al., 2007). In contrast, tissue-specific factors such as MEF2C and KLF in the cortex, and AR in the SCN showed decreased co-occurrence with remodelling factors. This finding re-emphasizes different roles of key sleep-responsive TFs and lineage-specific TFs that both regulate target gene expression in response to SD via accessible chromatin.

### The Contribution of Enhancer RNAs to the regulation of gene expression in specific **brain regions and in response to SD**

To identify actively transcribing enhancers (eRNAs) that could regulate brain region specific gene expression and the response to SD, we combined the outputs from the ATAC-Seq and directional bulk RNA-Seq datasets. Recently, eRNA has been shown to modulate tissue-and-stimulus (chemical and electrical) specific gene expression in highly co-ordinated manner to control the target mRNA levels (Joo et al., 2016; Kim et al., 2010; Telese et al., 2015). Generally, these non-coding RNAs are neither spliced nor polyadenylated (Arner et al., 2015; Gray et al., 2015) and are believed to aid long distance enhancer–promoter interaction to control target gene expression. Furthermore, incidences of circadian eRNAs, oscillating and peaking with distinct phases, have been observed in the mouse central clock (Bafna et al., 2023) and peripheral tissues such as liver (Fang et al., 2014) to bring about rhythmic gene expression. Therefore, using the directionality of the RNA-seq readouts, we identified intergenic OCRs that also displayed bi-directional transcription of RNA. These RNAs were classified as enhancer RNAs (Fig. 7a) (Arner et al., 2015; Barshad et al., 2023).

**Figure 7.**
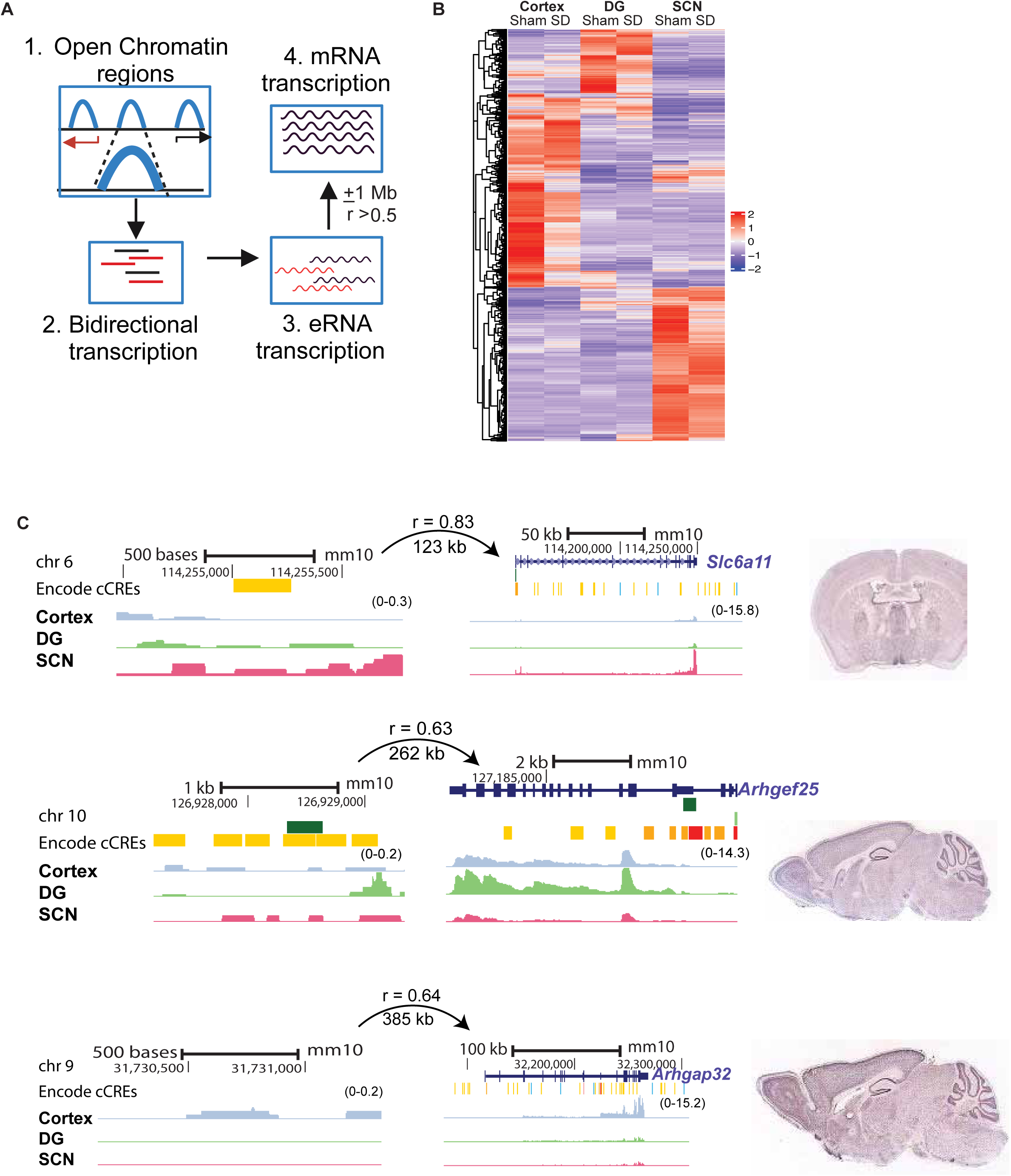
Region-specific enhancer RNAs (eRNAs) arising from open chromatin regions. *(A)* Schematic for identification of actively transcribing enhancers and determination of eRNA–gene pairs by correlation of eRNA and gene expression levels within a 1-Mb distance cutoff. *(B)* Clustered Heat map view of differential eRNAs observed selectively in CTX, DG and SCN*. (C)* UCSC Genome Browser tracks showing normalized eRNA and mRNA levels for specific examples from the SCN, DG and cortex. The distance between eRNA and predicted target gene with Pearson’s correlation is indicated above each represented eRNA–gene pair. Tissue-specific expression of the respective target gene is seen from *in situ* hybridization images from adult mouse brain (right; image credit: Allen Brain Atlas, Allen Institute).

In total, we identified 18,719 open chromatin regions (OCRs) across the three brain regions that also showed bi-directional transcription. Of the 18,719 identified putatively active enhancers (TAPEs), we observed 811 CTX, 286 DG and 722 SCN region-specific TAPEs (Fig. 7b, Supplementary Table 5). To detect the target mRNAs under the control of eRNAs, we examined the expression level of genes ± 1 mb from the site of bidirectional transcription (eRNA). We adopted Pearson’s correlation to study the association between the eRNA and target genes and assigned those eRNA-mRNA pairs with a positive correlation (coefficient ≥ 0.5) as being co-regulated. This strategy, based on expression abundance, has been previously reported to identify target genes that are under the control of transcribing enhancers (Carullo et al., 2020). We identified 22,915 positively correlated eRNA-mRNA pairs with global correlation > 0.5 (Supplementary Table 6). As expected, region-specific eRNA (Fig. 7b) showed strong association with region-specific gene expression (Fig. 7c). These included GABA transporter *Slc6a11* (Solute Carrier family 6 member 11) in the SCN, *Arhgef25* (Rho guanine nucleotide exchange factor 25 ) in DG, and *Arhgap32* (Rho GTPase activating protein 32) in CTX involved in protein sorting and trafficking in neurons (Teasdale & Collins, 2012) . We then conducted differential expression analysis to determine the effect of sleep deprivation on eRNA levels in each brain region. As reflected by the overall transcriptomic response to SD, we found a greater number of SD-regulated enhancer RNAs in the DG (n = 39) and CTX (n = 33), when compared to the SCN (n = 20) (Supplementary Table 7, Supplementary Fig. 4). Of these, only 8 eRNAs changed in both DG and CTX, again reflecting that the majority of changes in response to SD occur in a brain region-specific manner. Using the same approach, each differential eRNA was assigned to potential target mRNAs based on co-variation for each tissue. By doing so, we noticed that tissue-specific SD affected genes such as *Fosl1, Kcns2* in cortex, *Pcdhb21* in the DG and *Errfi1*in the SCN were under the influence of distal transcribed eRNAs.

### Effect of DNA methylation on sleep-deprivation

DNA methylation regulates target gene expression in a wide range of organisms in multiple ways. Promoter hypo-methylation and gene body hyper-methylation are associated with increased gene expression (Ball et al., 2009) and occurs on very long (generations) (Gruzieva et al., 2019; Lowe et al., 2018) as well as short timescales (Elbere et al., 2018; Glahn et al., 2014; Luo et al., 2018; Xie et al., 2023). To understand the contribution of DNA methylation to the observed changes in OCR and gene expression, we conducted reduced representation bisulfite sequencing (RRBS) (Beck et al., 2022; Huang et al., 2022; Nakabayashi et al., 2023; Simpson et al., 2023) using CTX and SCN tissues from sham control and SD mice. We assessed genome-wide DNA methylation patterns and annotated differentially methylated regions (DMRs) by comparing a sliding window of 10 differentially methylated sites across brain region and/or condition. Specifically, we noted 661 DMRs between CTX and the SCN (Fig. 8a), with the majority concentrated around the promoter region (Fig. 8b). We then inspected the intersection between differentially expressed genes (CTX vs SCN) and tissue-specific methylation. Surprisingly, we found that only 3.49% of the differential genes (80 out of 2289) showed significant change in methylation levels, suggesting DNA methylation changes contribute a small but significant amount to the tissue specific gene expression. Next, we assessed the prevalence of DMRs between sham controls and SD mice in a tissue-specific manner. Remarkably, 6-h acute sleep perturbation resulted in significant difference in DNA methylation across the tested brain regions (Supplementary Table 8), with genes showing both hypo-and-hyper DNA methylation owing to SD (Fig. 8c, d). By this approach we successfully uncovered 289 and 377 DMRs between sham and SD conditions in the SCN and cortex respectively. For example, following SD decreased methylation was clear for the SCN-specific gene *Gnas (Brown et al., 2017)* near the gene TSS (Fig. 8e). This is in agreement with previous studies where *Gnas* has been linked to sleep such that loss of *Gnas* imprinting improved NREM, but worsened REM sleep *(Lassi et al., 2012; Tucci et al., 2019)*. Overall, our current dataset supports the idea that DNA methylation is highly dynamic and under tissue-and-context specific regulation. It is noteworthy that DNA methylation can also influence chromatin accessibility which can further regulate downstream gene expression (Chang et al., 2023; Zhong et al., 2021). This hierarchical regulatory mechanism is exemplified by gene *Map3k6* (mitogen-activated protein kinase 6, Supplementary Fig. 5), where intragenic hypomethylation is seen to be associated with increased chromatin accessibility facilitating enhanced gene expression under SD in the SCN. Taken together, our findings suggest that the transcriptional response to sleep deprivation is unique to each brain region, and this response is coordinated by a complex grammar encoded at the level of chromatin architecture, and the illustration of *Map3k6* provides an elegant visualisation of this regulatory grammar.

**Figure 8.**
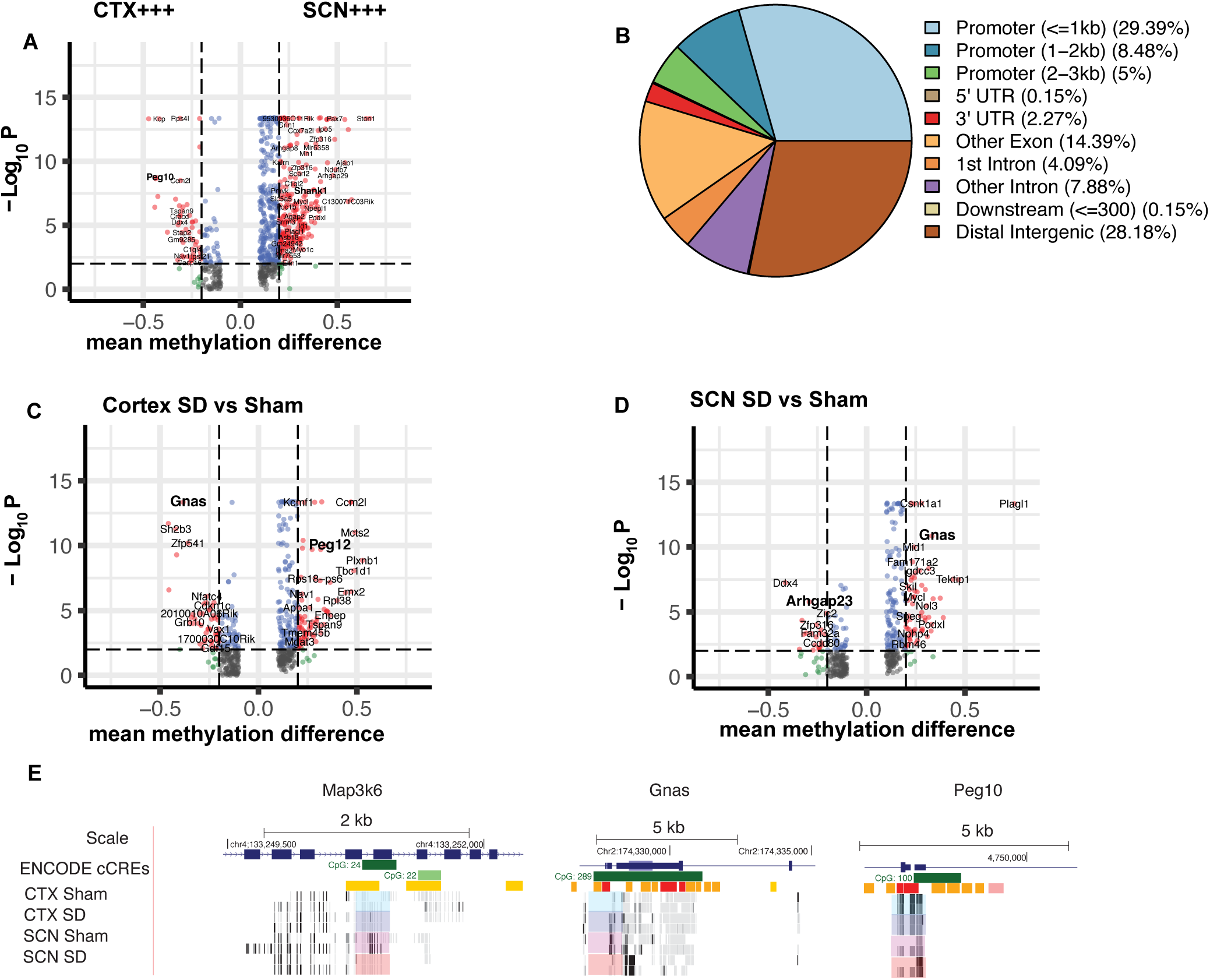
Differential CpG methylated regions (DMRs) as a key regulator of SD response. *(A)* Volcano plot showing mean methylation difference and logarithmic P-value (log_10_P) as computed by the tool Metilene using cutoff (d) at 0.1 between SCN and CTX. *(B)* Genomic feature distribution of differential DMRs between SCN and CTX as computed by ChIPSeeker. *(C)* Volcano plot showing mean methylation difference and logarithmic P-value (log_10_P) as computed by the tool Metilene using cutoff mean methylation difference at 0.1 between Sham and SD in CTX and *(D)* SCN. *(E)* UCSC Genome browser tracks of the gene loci *Map3k6*, *Gnas* and *Peg10* which display region and/or SD-response specificity in the prevalence of DMRs in CTX (blue) and the SCN (pink). The intensity of the filled bar is directly proportional to observed methylation fraction (black = 100% methylation, white = 0% methylation at the CpG site). The chromosome location and scale (mm10 genome) are indicated at the top along with ENCODE tracks for cCRE elements and CpG methylation.

## DISCUSSION

In this study we use the power of multi-omics study to understand how the transcriptomic response to sleep deprivation is dynamically regulated in distinct brain regions. Through the integrated analysis of genome-wide DNA methylation, chromatin accessibility and total RNA expression, we identified that the transcriptome changes in a tissue-and- SD dependent fashion, and that this response is coordinated at the level of upstream regulators that change chromatin accessibility, DNA methylation (CpG), TF binding and enhancer activity, in response to 6-hour sleep loss. To the best of our knowledge, the current study is the first of its kind to demonstrate that both enhancer activity (eRNAs) and DNA methylation change rapidly in response to SD in a tissue-specific manner. Together, these data provide a bird’s eye view of action within regulatory elements encrypted in the genome that enable different brain regions to mount a distinct response to in sleep loss in a manner consistent with their own physiological roles.

From the transcriptomic analysis, we noted that gene expression could accurately reflect brain region-specific functions (Fig.1), which differ considerably between cortex, DG and the SCN. Such a phenomenon has been described for various peripheral tissues (Kouadjo et al., 2007; Srivastava et al., 2020), but rarely in separate brain regions due to technical limitations. Whilst we observed several transcripts were affected by SD throughout the brain, (e.g *Cirbp, Dbp*) many showed distinct region-specific changes, supporting a unique role for each region in sleep regulation. For instance, differences in the neuropeptide expression levels were seen only in the central pacemaker, suggesting SD would affect intercellular coupling in the SCN alone, and not in the DG or CTX. This highlights the importance of examining separate brain regions to fully understand the orchestration of sleep regulation in mammals. A limitation of our study is that we conducted bulk sequencing, whereas specific combinations of TFs, OCRs and gene expression that co-occur within specific cell-types may be better captured using a single-cell multiomics approach, which to our knowledge, has not been attempted to date. Single-cell transcriptomics has highlighted the diversity in cell-type responses to SD (Dopp et al., 2024; Jha et al., 2022; Vanrobaeys et al., 2023). However, by comparing multiple omics readouts in different brain regions, we were able to correlate the genomic regulome to transcriptional level changes that were specific to each region in response to SD.

An important question is what determines such brain-region specific transcriptional networks. To identify the determinants of such differences in transcriptional regulation, we profiled dynamic chromatin architecture. We confirmed previous findings that SD causes widespread changes in chromatin accessibility (Hor et al., 2019), but saw that the patterns of change depended on the brain region sampled. We saw that open chromatin regions (OCRs) correlated with target gene expression in multiple ways-(1) A direct correlation as exemplified by *Hif3a* gene (Fig. 4), where increased promoter accessibility accompanied elevated gene expression under SD. (2) An inverse correlation as seen with *Cirbp,* suggesting the binding of repressive factors at the OCR. 3) No change in chromatin accessibility in relation to the gene expression levels. 4) Change in chromatin accessibility at a distal site target gene expression, as seen for *Homer1* (in CTX) and *Gabre* (SCN). Moreover, in some cases multiple open chromatin regions (OCRs) with different responses to SD were found proximal to the same target gene, indicating a co-operative and non-linear pattern of gene regulation. An important outcome from our analysis of these different patterns is that it is not meaningful to assign one or many OCRs to genes on the basis of the nearest neighbour approach (Chen et al., 2017; Kim et al., 2010). This could be why chromatin accessibility (represented by individual genomic sites) was not seen as a reliable predictor of the gene transcription, as suggested by Carullo et al (Carullo et al., 2020). Rather, chromatin accessibility reflects the sum of a complex series of modifications which regulate transcription in different ways, such as DNA methylation, histone modification and the binding of both activating and inhibiting transcription factors at promoters and enhancers. Therefore, we undertook a number of follow-up approaches – both genome-wide as well as fine-grained, to identify the putative mediators of chromatin accessibility changes on a case-by-case basis.

Systematic genome-wide footprinting analysis identified key TFs determining both region specificity and the response to SD. Crucially, many of the tissue-specific TFs were also responsive to SD, such as AR in the SCN, MEF2C in cortex, and the binding of a vast majority of such region-specific TFs was reduced after SD. For example, the binding of CLOCK::BMAL1, encoded by the *Arntl* gene, was reduced in CTX after acute sleep loss corroborating previous findings by Mongrain et al. (Mongrain et al., 2011). Similarly, the nuclear mineralocorticoid receptor NR3C2 and AR (androgen receptor) were depleted in the SCN after SD relative to sham conditions, suggesting a role for tissue-specific TFs in shaping the regional responses to SD (Fig. 6b). AR is expressed mainly in the retinorecipient SCN cells and administering androgen *in vivo* has been shown to enhance light -induced phase shifts of the circadian clock (Butler et al., 2012). On the other hand, SD is known to attenuate photic phase-shifts (Mistlberger et al., 1997) and the data from our footprinting analysis suggesting reduced AR binding following SD could underlie the mechanism via altering the chromatin landscape of the SCN that “presents” to light. Furthermore, in all brain regions, the co-occurrence of chromatin / DNA methylation remodellers such as ARID3b and DNMT3a (DNA methyltransferase 3A) with tissue-specific TFs was reduced after sleep loss. Interestingly, these remodellers now co-occurred preferentially with AP-1 related factors. AP-1 is a family of over 20 transcription factors (FOS, JUN, MAF and ATF families) that are involved in nearly every physiological process including sleep and circadian rhythms (Jagannath et al., 2021). As an activity-dependent TF family, the AP-1 members including FOS have been shown to have a critical role in initiating chromatin opening and thus remodelling the genomic landscape in a manner that reflects environmental signals or activity (Su et al., 2017; Vierbuchen et al., 2017). In the case of SD, whilst FOS has long been studied as a marker of sleep loss (Cirelli 1995), its role has remained somewhat obscure. Our data show that AP-1 binding increases in all tissues after SD, but that regulatory elements in the genome on which these factors act are shaped by the repertoire of tissue-specific factors available. Furthermore, the diversity of AP-1 factors and their binding sites could explain the differential responses to SD (upregulation of some genes, downregulation of others) despite an increase in AP-1 binding. For example, and as described above, a binding site for the AP-1 repressor element BACH1 was found in the promoter region of *Cirbp,* which could explain its reduced expression after SD despite increased chromatin accessibility at the promoter following SD. Thus, the results of the TF footprinting analysis of OCRs show that there are distinct region- and SD-specific transcription factors that interact to shape the dynamic chromatin accessibility landscape to modulate target gene expression in response to SD.

Enhancers are known to regulate target genes in a context and stimulus (electrical, chemical, environmental) dependent fashion. Enhancer activity can occur at remarkably short timescales that are relevant to neuronal activity - Beagan et al. demonstrated the formation of enhancer-promoter loops within 20 min of stimulation at IEGs including the *Fos* gene (Beagan et al., 2020). We used the power of combining chromatin accessibility changes with concomitant expression of non-coding RNAs to identify active enhancers and found a repertoire of region-specific and/or SD responsive eRNAs. Interestingly, with this approach, we also noted that a single OCR could elicit bidirectional eRNA transcription that could further control one or many genes, based on co-variation (Supplementary Table 8). These findings extend our existing knowledge on the regulation of genes in the context of sleep. For example, we found an eRNA at chromosome 19 (Chr. 19: 5821709-5823003) potentially regulating enhancer-promoter loop formation and therefore the expression levels of *Fos* gene in the cortex (r = 0.77) in response to SD. In future, a detailed investigation of enhancer-promoter contact tracing by means of 3-C approach will validate the mapped long-range interactions. It will be highly beneficial to further investigate the effect of enriched region-selective eRNAs by means of targeted mutagenesis and study their role in mediating enhancer-promoter interactions that are essential to potential target gene(s) to gain mechanistic insights into sleep regulation. Nonetheless, we clearly catalogue region-and-sleep select eRNAs that potentially refines the enhancer-promoter interactions which are essential to generate lineage and/or context dependent response.

Lastly, we addressed the role of DNA methylation by examining differential CpG methylation. Whilst we identified hundreds of brain region-specific methylation sites, we also noted changes in DNA methylation at relatively short timescales - 6h of sleep deprivation - and their functional association with changing chromatin accessibility and transcription (as exemplified by *Map3k6)*. These data provide substantial evidence that DNA methylation is an essential part of the repertoire alters chromatin accessibility and regulate gene transcription (Mansisidor & Risca, 2022) in response to sleep loss.

Furthermore, for certain genes such as *Gnas*, DNA methylation changed in response to SD in both cortex and the SCN, but at distinct genomic sites. As illustrated in Supplementary Fig. 6, we noted three distinct DMRs (Sham vs SD) associated with cortex, while only one changed in the SCN, intriguingly within the CpG hotspots (ENCODE) next to different transcript variants of the gene. In addition, the methylation level decreased in response to SD in the SCN, but increased in the CTX, reflecting the diverse roles of DNA methylation in tissue-and-condition specific gene regulation.

In summary, our data provide new insight into the gene regulatory networks that allow a generic stimulus such as sleep deprivation to elicit distinct molecular and physiological outcomes in different regions of the mammalian brain. Overall, we show how multiple elements of gene regulatory network converge into a complex and fine-tuned system to relate to whole brain physiology. Furthermore, our datasets provide a valuable resource to explore multi-faceted gene regulatory networks in a functional context.

## METHODS

### Experimental Subjects and Models

All studies were conducted on C57Bl/6J mice over 50 days of age of both sexes. Unless otherwise indicated, animals were group housed with food and water *ad libitum* under a 12:12 h LD cycle. All procedures were performed in accordance with the UK Home Office Animals (Scientific Procedures) Act 1986 and the University of Oxford’s Policy on the Use of Animals in Scientific Research (PPL 70/6382, PPL 8092CED3), as approved by the local Animal Care and Ethical Review committee (ACER). Animals were sacrificed via Schedule 1 methods in accordance with the UK Home Office Animals (Scientific Procedures) Act 1986.

### Experimental design

C57Bl/6 mice were housed under a 12:12 LD cycle for 2 weeks with food and water *ad libitum*. The sleep deprivation protocol consisted of novel object introduction and manual gentle handling between ZT 0-6. Animals were sacrificed by cervical dislocation and brains were removed from sham treated and SD mice, and placed into a brain matrix (Kent Scientific, Torrington CT, USA). A skin graft blade (Swann-Morton, Sheffield, UK) was positioned at Bregma −0.10 mm. A second blade was placed 1 mm caudal from the first, and a 1 mm thick brain slice was dissected. Cortex, DG and the SCN punches were taken using a sample corer (1 mm internal diameter, Fine Science Tools GmbH, Heidelberg, Germany) from the brain slice, flash frozen on dry ice and stored at −80 °C prior to genomic DNA/RNA extraction. For ATAC-seq, n = 2 per tissue (cortex, DG, SCN)/ condition (sham or SD) was used. For RNA-seq, n = 8 per tissue/condition was used. For RRBS sequencing, n = 6 per tissue (cortex, SCN)/ condition (sham or SD) was used for library preparation.

### ATAC-Seq library preparation and sequencing

Tissue punches (cortex, DG, SCN) were homogenized directly to nuclei by adopting the nuclei isolation protocol from Su et al. (Su et al., 2017) with slight modifications (using a dounce homogenizer and sucrose cushion). The viability of nuclei was then checked under a microscope using ThermoFisher NucBlue™ Live ReadyProbes™ reagent. Following successful isolation of intact nuclei, ATAC-seq libraries were prepared using the Tn5 transposase. Typically, nuclei from 4 punches, containing approximately 40,000 nuclei were re-suspended in the transposition mix (25μL 2x TD Buffer (Illumina Cat no. FC-121-1030), 2.5μL Tn5 Transposes (Illumina Cat no. FC-121-1030) and 22.5μL nuclease free water) and incubated at 37°C for 30 min. DNA was then purified using the Qiagen MinElute Kit and eluted with 23μL warm 10mM Tris, pH8 and stored at -20°C. The transposed DNA was finally amplified using Nextera sequencing primers, quantified and sequenced at Illumina NextSeq 500 platform.

### ATAC-Seq mapping and peak calling

Quality assessment of paired-end FASTQ files was performed using FastQC v0.11.9 (Wingett & Andrews, 2018), and adapters were trimmed using Trim Galore v0.6.2 https://www.bioinformatics.babraham.ac.uk/projects/trim_galore (parameters: --phred33 --quality 15 --stringency 1 -e 0.1 --length 20). Trimmed sequences were aligned to the mouse mm10 genome with Bowtie2 v2.3.5.1 (parameters: --end-to-end --very-sensitive --phred33 -X 2000) (Liu & Schmidt, 2012). Reads aligning to the mitochondrial chromosome (chrM) were filtered using Samtools v1.8 (Danecek et al., 2021). PCR duplicate reads were marked using Picard v2.21.6 (Broad Institute. (Accessed: 2018/02/21; version 2.17.8) http://broadinstitute.github.io/picard/. Unmapped, duplicated, and low-quality reads were then removed by Samtools using the –F 1548 flag. Multi-mapped reads were removed by retaining only reads with MAPQ score > 30. Samtools, was then used to sort and index the filtered alignment files. Blacklist regions were filtered using Bedtools v2.29.2 (Quinlan & Hall, 2010), using the blacklist database from ENCODE (Li et al., 2009). Filtered binary alignment map (BAM) files for technical replicates were merged using Samtools. MACS2 -defined ATAC-seq peak calling with the options --qvalue 0.00001 --gsize 2652783500 --format BAMPE was performed for each brain region, and peaks closer than 1000 bp were merged with BEDtools. The resulting region specific ATAC-seq peaks were combined to generate a final list of 150,234 regions of open chromatin (OCR). For the differential peak analysis Diffbind (https://bioconductor.org/packages/release/bioc/html/DiffBind.html) was executed at FDR <0.05 using summits ± 100 bp of the identified peaks in tissue and/or condition dependent way. Differential peaks were assigned to closest TSS (± 3kb) using ChIPSeeker v1.28.3 (Yu et al., 2015) and reported with distance and strand information.

### Enhancer RNA identification

Enhancer identification was performed on merged ATAC-seq and total RNA-seq BAM files for Sham and acute sleep-deprived (SD) samples from cortex, dentate gyrus (DG) and suprachiasmatic nuclei (SCN) tissues using the published pipeline (Carullo et al., 2020). Briefly, MACS2 -defined ATAC-seq peak calling with the options --qvalue 0.00001 --gsize 2652783500 --format BAMPE was performed for each brain region, and peaks closer than 1000 bp were merged with BEDtools. The resulting region specific ATAC-seq peaks were combined to generate a final list of 150,234 regions of open chromatin (OCR) spanning ± 500 bp of identified ATAC-seq peaks. We then filtered out OCRs that overlapped with known noncoding RNAs such as miRNA, rRNA, snoRNA, snRNA and tRNA using Seqmonk https://www.bioinformatics.babraham.ac.uk/projects/seqmonk/loaded_with_genomeGRCm38_v100. These filtering steps provided a list of 45,245 intergenic OCRs (iOCRs).

Bidirectional transcription at iOCRs was calculated in Seqmonk, and a list of 18,719 transcriptionally active putative enhancers (TAPEs) was compiled. Possible TAPE-gene pairs were identified by mapping all gene promoters within 1Mb upstream or downstream from the centre of the TAPE. CPKM values for each TAPE and associated gene were correlated using a Pearson’s correlation in R (https://www.R-project.org/). Pearson’s correlations were calculated using all samples (to give a global correlation across tissues), as well as using region-specific samples (to generate region-specific correlation values).

TAPE–gene pairs with correlations > 0.5 were deemed high-confidence pairs. These high-confidence pairs (n = 22,915) were then used to investigate distance and gene position distributions. Region-specific TAPEs were identified in Seqmonk using DESeq2 (Love et al., 2014) to compare TAPE RNA counts in cortex, dentate gyrus and the SCN (FDR < 0.05) using all replicates for Sham controlled samples. DESeq2 identified 1819 potential region-selective TAPEs. However, this list includes TAPEs which may be highly expressed in two of the three culture systems. To ensure region selectivity, we conducted hierarchical clustering on TAPE RNA values from these 1819 TAPEs (z-scored across region) in Seqmonk to identify nodes of TAPEs specific to each region. This analysis identified 811 cortex, 286 DG and 722 SCN region-selective TAPEs respectively.

### Footprinting Analysis

The list of mouse transcription factor (TF) motifs was sourced from (Yi et al., 2021), which was compiled from existing mouse TF binding site databases CIS-BP 1.02 (Weirauch et al., 2014), JASPAR 2022 (Castro-Mondragon et al., 2022), and HOCOMOCO v11 (Kulakovskiy et al., 2018). At most three motifs per gene were kept, resulting in a total of 1636 motifs from 836 mouse transcriptional regulators. For comparisons between SD and sham in each brain region, following filtering steps to remove mitochondrial reads, blacklist regions, and PCR duplicates, plus down-sampling to account for library size differences between technical replicates in the same brain region, BAM files for each brain region (cortex, dentate gyrus, and suprachiasmatic nuclei) were merged. To increase resolution for the TF footprinting analysis, summits were called during MACS2 peak calling using the options --nomodel --shift -73 --extsize 146 -g 2407883318 --keep-dup all --call-summits --nolambda -p 0.01. Summits were then extended by 250 bp to both sides to a fixed width of 501 bp. Peaks that extend beyond chromosome ends were filtered. The remaining peaks went through an iterative removal process to keep the most significant peaks and remove less significant peaks that overlap with more significant ones, as described by (Corces et al., 2018). The result is a region-specific peak set for each brain regions. For comparisons across brain regions, a similar procedure was undertaken, except some adjustments mentioned below. BAM files of technical replicates in each brain region were merged, based on which peak calling was performed to generate a consensus peak set for each brain region. To normalize the differences between brain regions in terms of read depth and quality, individual MACS2 501-bp peak score (“-log10(pvalue)”) was divided by the sum of all peak scores in each sample and multiplied by 1 million. The peak sets for each brain region were then merged using Bedops v2.4.36 (Neph et al., 2012) and were iteratively filtered again to retain significant peaks from all brain regions to generate a master peak set representing producible peaks from all brain regions. This set can then be used for comparisons across brain regions. Assessment of TF footprints were assessed using the TOBIAS pipeline v0.2.10 (Bentsen et al., 2020). Briefly, using processed ATAC-seq peak sets and list of known mouse TF motifs, the degree of TF binding across both conditions (sham vs SD) and across brain regions was assessed. ATACCorrect tool was used to correct the Tn5 Transposase insertion bias, and then a continuous footprinting score was calculated from the bias-corrected cut sites, considering signal depletion and genome accessibility of nearby regions. Footprint scores across conditions and brain regions were then compared to identify differential footprint regions, corresponding to differential binding of specific TFs.

### RNA-Seq library preparation and sequencing

100 ng of RNA was used to prepare libraries using the Illumina Stranded Total RNA Prep, Ligation with Ribo-Zero Plus kit following the manufacturer’s instructions. Following library preparation, library concentration was determined using the Library Quantification Kit for Illumina Platforms (Roche, UK) following the manufacturer’s instructions. RNA-seq was performed as 150-bp paired-end sequencing on Illumina platform by Novogene (Cambridge, UK).

### RNA-Seq mapping and analysis

The quality of raw sequencing data was assessed using FastQC v0.11.9 (Wingett & Andrews, 2018) both before and after adapter trimming. The reads were trimmed for Nextera adapter sequences, low-quality base calls (Phred score < 15) and short reads (read length < 20 base pairs) using Trim Galore v0.6.2. The processed FASTQ paired-end reads were aligned using HISAT2 v2.2.1 (Kim et al., 2019) to mouse GRCm38 genome build from Ensembl. BAM files were sorted by read name and chromosome position using Samtools v1.8 (Li et al., 2009). Transcripts were quantified via the FeatureCounts function of Rsubread v1.6.4 (Liao et al., 2019). For comparison between different brain regions and/or condition (sham vs SD), downstream differential analysis was performed on the count matrix using R version 4.2 and R package DESeq2 v3.15 (Love et al., 2014). Data was normalized using DESeq2’s built-in median-of-ratios method to account for library depth and RNA composition across samples. Genes with low counts (sum of counts is less than 30 across all samples) were filtered, as they mostly reflect noise in the dataset. Genes that are differentially regulated by SD in a brain region-specific manner were computed at p-adjusted < 0.1 in the region of interest with p-adjusted > 0.6 in other regions. Heatmaps were generated by with the R package ComplexHeatmap (v2.18.0, available at: DOI: <u>10.18129/B9.bioc.ComplexHeatmap</u>) and volcano plots with EnhancedVolcano (v1.20.0, available at DOI: <u>10.18129/B9.bioc.EnhancedVolcano</u>)/

### RRBS library preparation and sequencing

Genomic DNA was extracted from the SCN and cortex of the sham control and sleep-deprived mice using MagAttract HMW DNA kit (Qiagen). 100 ng of the extracted DNA was used per tissue and condition to prepare RRBS libraries using Premium RRBS kit V2x96 for low DNA amounts and accurate analysis (https://www.diagenode.com/en/p/premium-rrbs-kit-V2-x24). The prepared libraries were sequenced (paired-end directional reads) on a Illumina-HiSeq platform.

### RRBS mapping and peak calling

Quality assessment of paired-end FASTQ files was performed using FastQC v0.11.9 and adapters were trimmed using Trim Galore v0.6.2. Trimmed sequences were imported at Galaxy web platform (Afgan et al., 2018) and aligned to the mouse mm10 genome with bwa-meth (Sun et al., 2018) v0.2.6 using paired-end option. Methylation calling was performed on aligned reads with Methyldackel v0.3.0.1 using the mm10 genome with options --mergeContext --fraction (https://github.com/dpryan79/MethylDackel.git).

Methylation scores were visualized with the UCSC Genome Browser (Kent et al., 2002). Differentially methylated regions (DMRs) were determined by Metilene v0.2.6.1 (Juhling et al., 2016) and defined based on --maxdist 300 --mincpgs 10 --minMethDiff 0.1 --mode 1 --valley 0.7 with p values <0.05 (as determined by MWU test). The nearest (± 3kb) gene to the identified DMRs was annotated by a Bioconductor package ChIPSeeker v1.28.3 (Yu et al., 2015) and reported with distance and strand information to the closest TSS (transcription start sites).

### Gene annotation

The gene list derived from DeSeq2 was fed into the gProfiler (version: e111_eg58_p18_30541362) (https://biit.cs.ut.ee/gprofiler/) as ENSEMBL Gene IDs. The functional annotation chart based on KEGG pathway and Gene Ontology was plotted with the heatmap for the selected terms.

## ACKNOWLEDGEMENTS

This work was supported by the following sources of funding: BB/N01992X/1 David Phillips fellowship BB/N01992X/1 from UKRI-BBSRC to AJ, and Oxford-Elysium Cellular Health Fellowship to LT and AB.

## AUTHOR CONTRIBUTIONS

AJ designed and supervised the study. LT and QD conducted all animal experiments, sample collection and processing. AB, QD, LT, CG, AJ and SW conducted the formal data analysis. Data visualisation was performed by AB, CG, DS and AJ. The original draft was written by AB and AJ. Manuscript was reviewed, edited and finalised by AB, QD, CG, DS and AJ.

## DATA AVAILIBILITY

All raw and processed sequencing data generated in this study will be submitted to the NCBI SRA upon acceptance.

## Figure Legends

**Supplementary figure 1.**
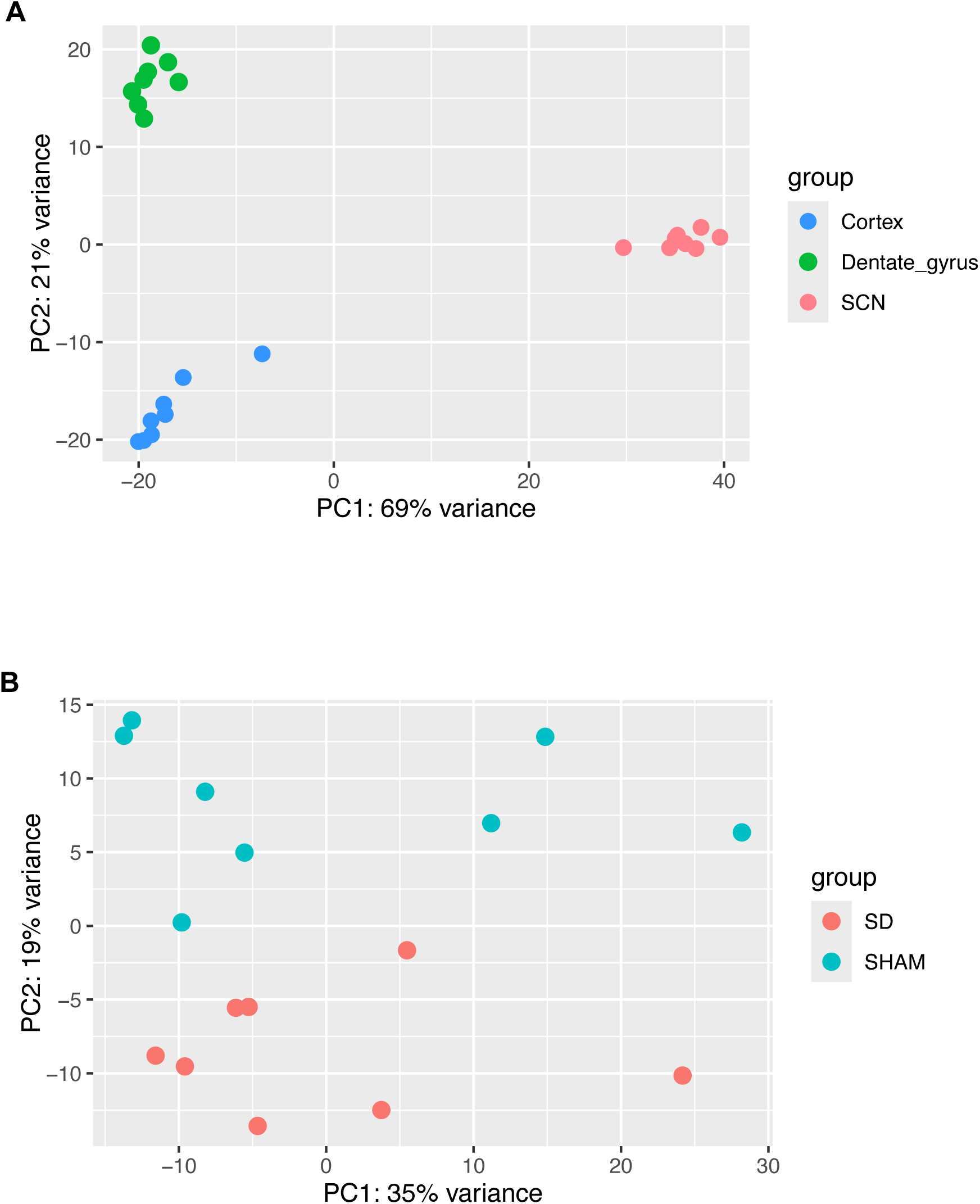
Principal component analysis (PCA) of RNA sequencing. *(A)*PCA analysis of normalised aligned RNA sequencing reads from CTX, DG and SCN SHAM (n =8/region) samples. *(B)* PCA analysis of normalised aligned RNA sequencing reads from SHAM and SD (n =8/condition) samples in CTX.

**Supplementary figure 2.**
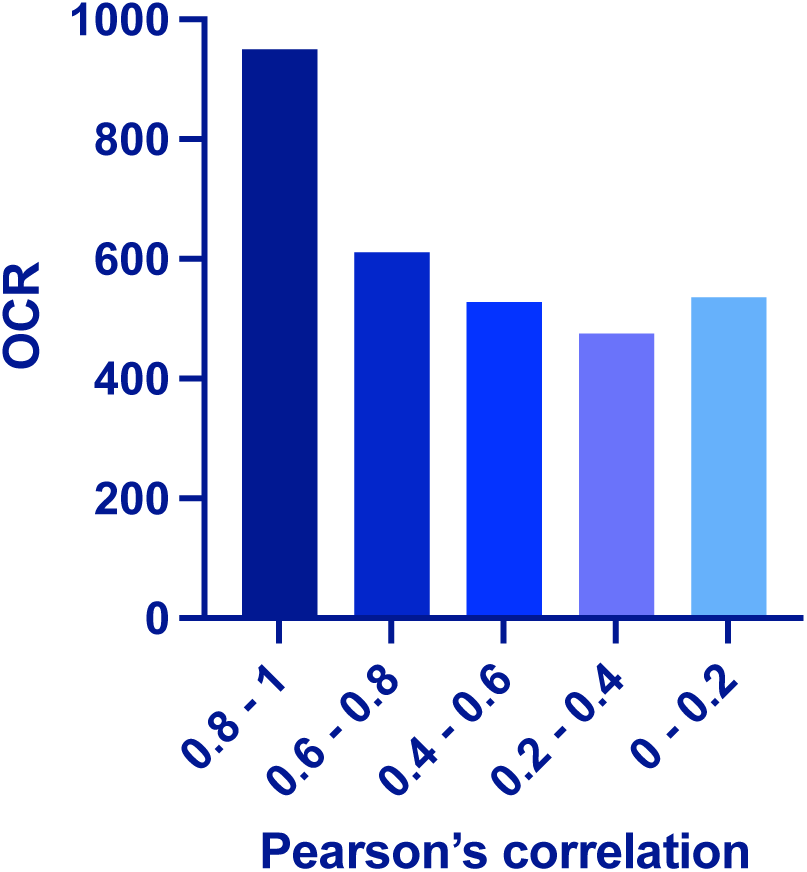
Strength of correlation between regional open chromatin regions (OCRs) and closest gene (TSS) under sham conditions. Bar-plot showing number of OCRs on y-axis and Pearson’s correlation between accessible chromatin and tissue-specific gene expression under sham conditions.

**Supplementary figure 3.**
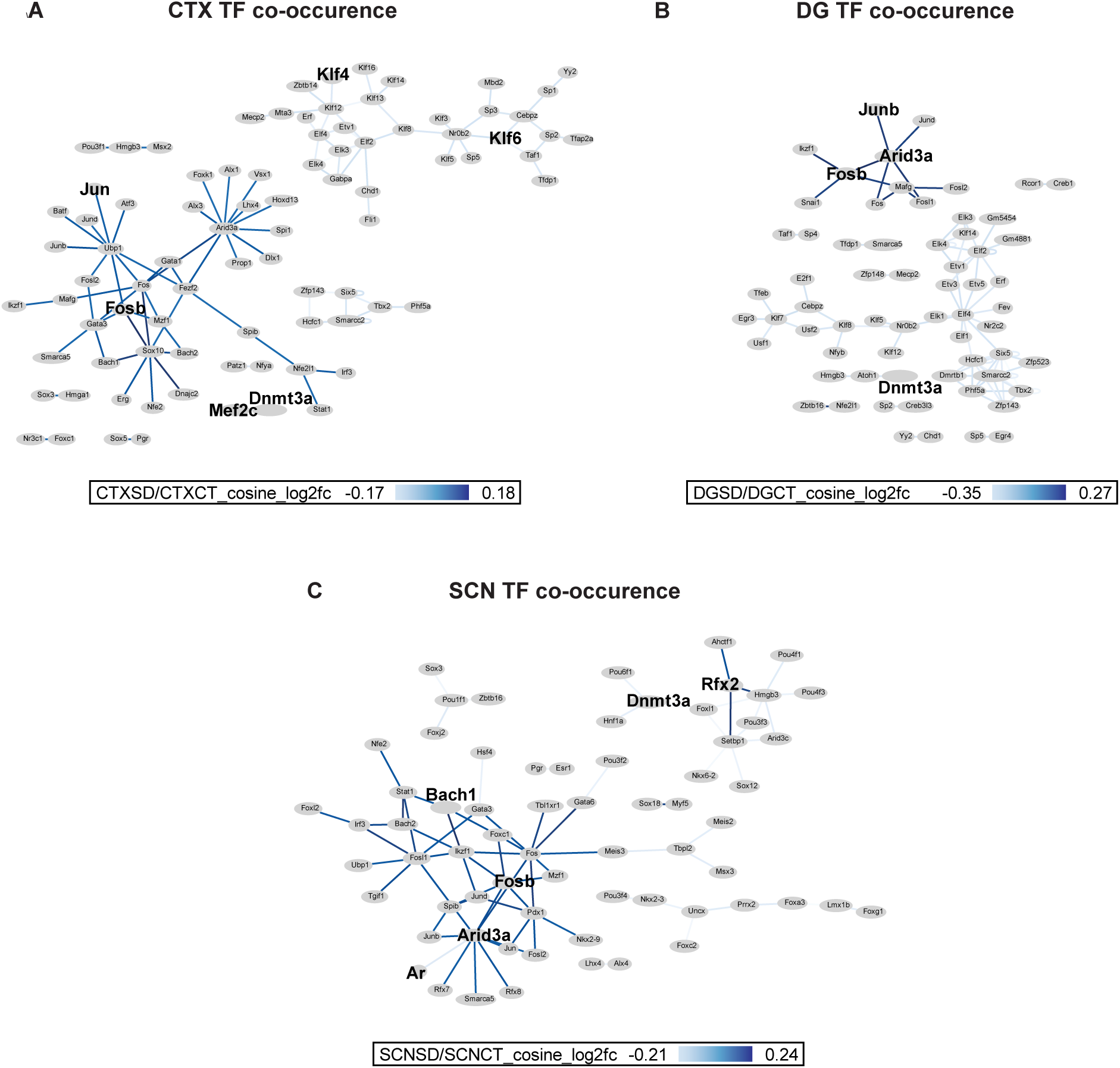
Co-occurrence of TFBS in SD response. *(A,B,C)* Transcription factor interaction plots for cortex (A), DG(B) and the SCN(C) in Sham vs. SD conditions. TF with greater interaction density in the SD are highlighted. The scale of interaction is shown below each plot.

**Supplementary figure 4.**
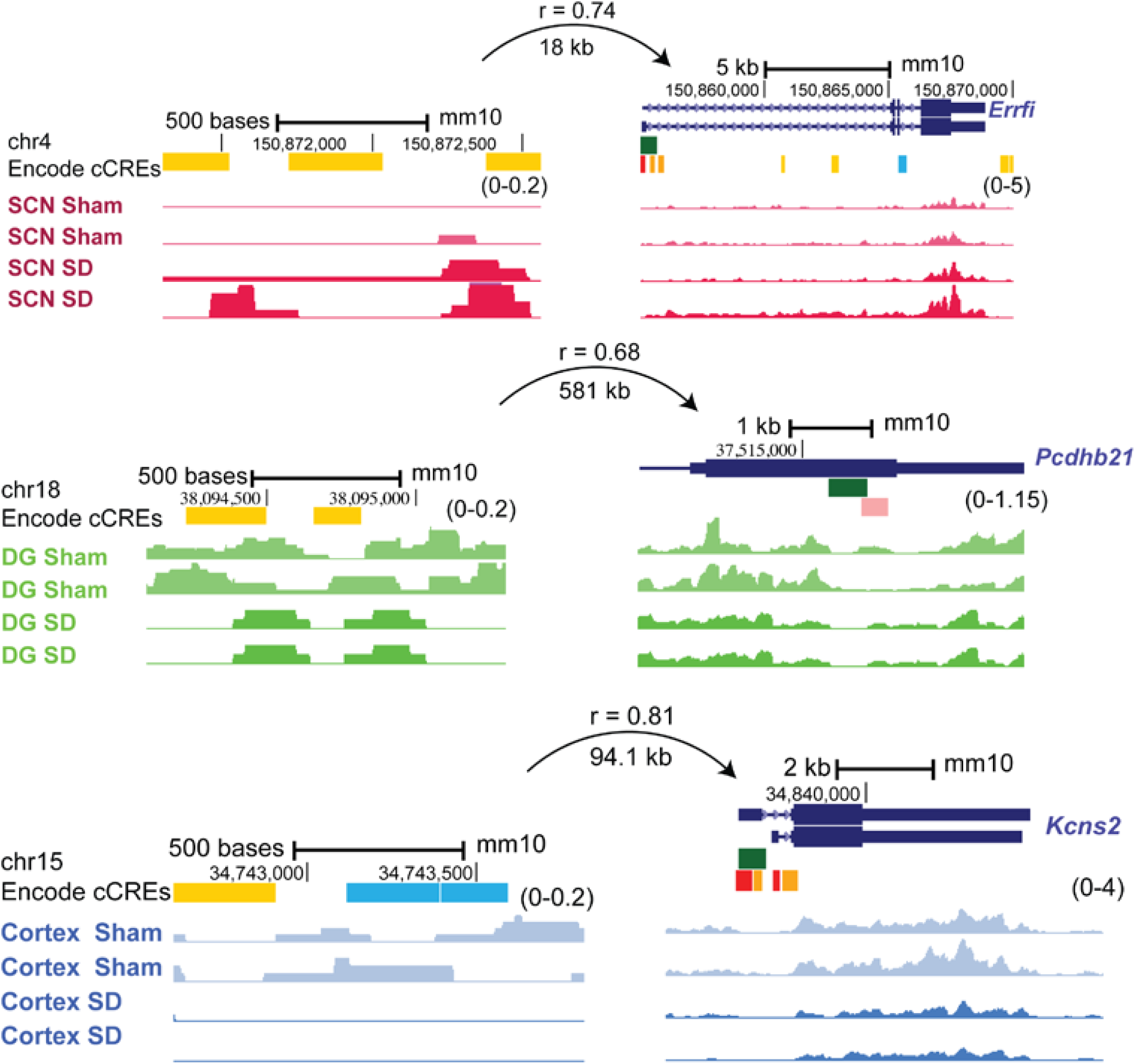
Positive correlation between the eRNA and mRNA in sleep-deprivation. UCSC Genome Browser tracks showing normalized eRNA (left) and mRNA expression levels (right) for specific examples from the SCN (pink), DG (green) and cortex(blue) regions showing differential expression between sham vs SD conditions. The distance between eRNA and predicted target gene with Pearson’s correlation is indicated above each represented eRNA–gene pair.

**Supplementary figure 5.**
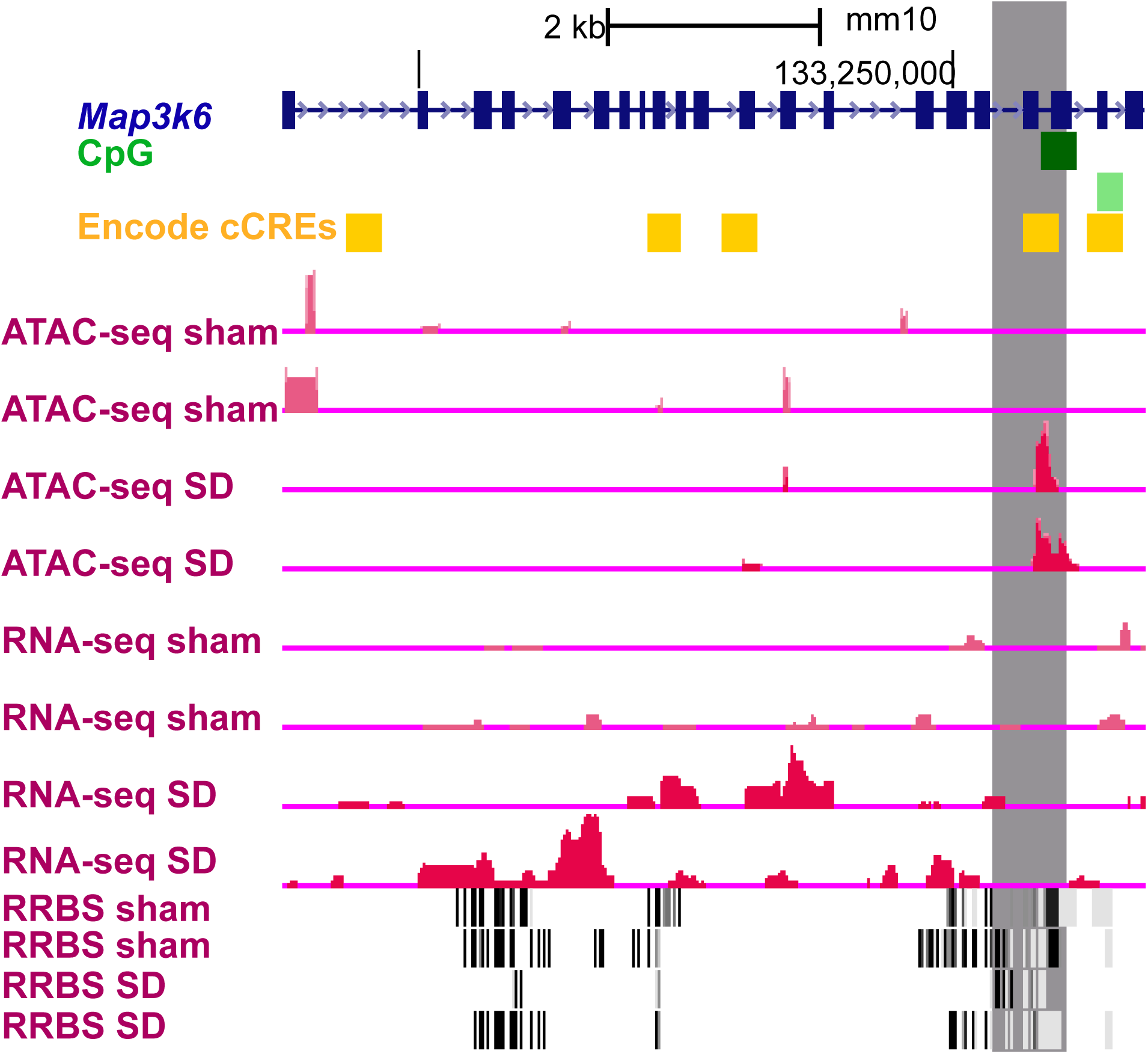
Representative example of correlation between the ATAC-seq, RNA-seq and RRBS datasets. UCSC Genome Browser tracks showing normalized ATAC-seq coverage, RNA-seq and RRBS for the gene *Map3k6*. The grey shaded region denotes differential chromatin accessibility (padj < 0.02), methylation region (p < 0.0002) and RNA levels (padj < 0.0005) between sham and SD in the SCN. The chromosome location and scale (mm10 genome) are indicated at the top along with ENCODE tracks for cCRE elements and CpG methylation.

**Supplementary figure 6.**
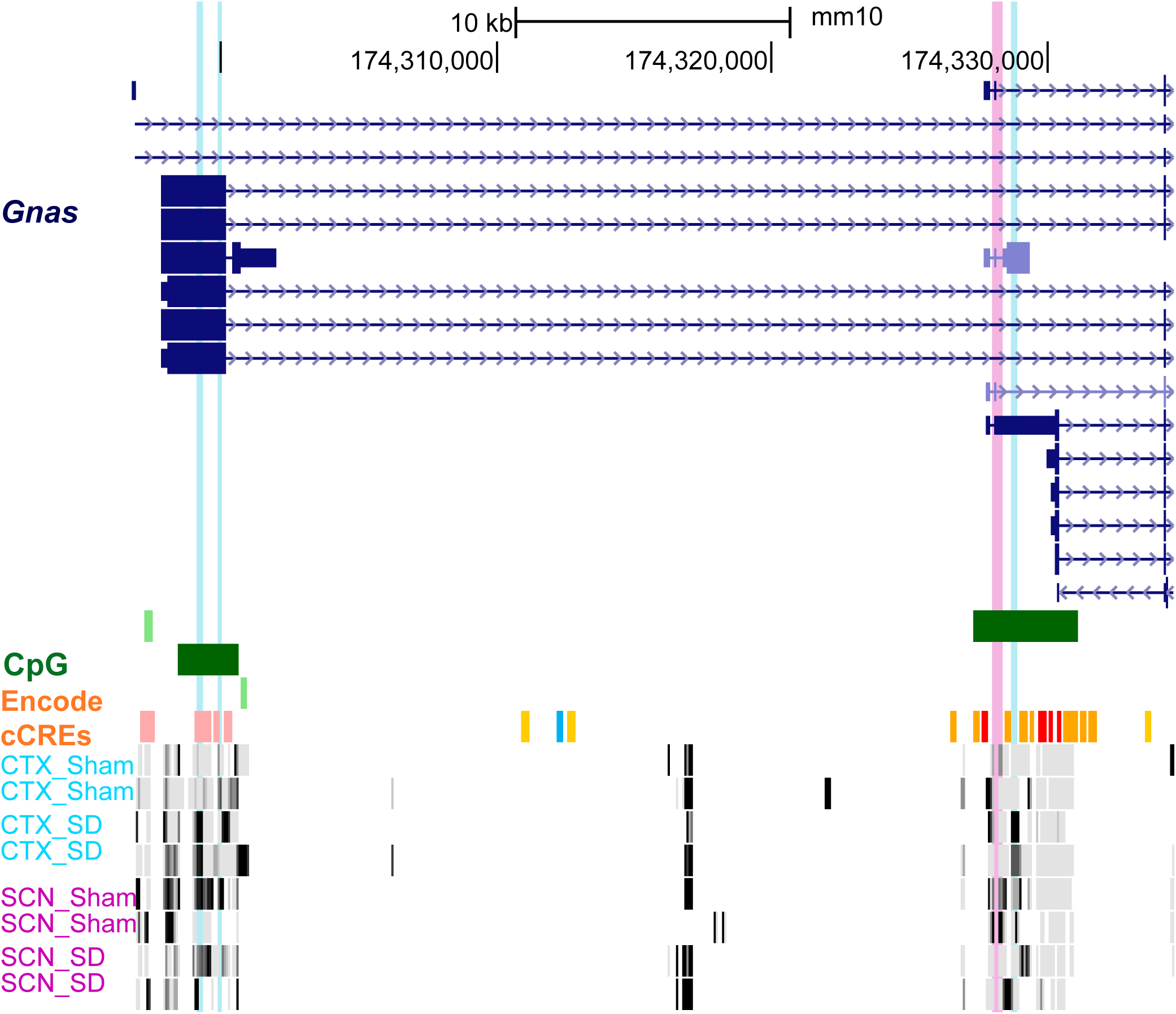
Representative example of differential methylation regions in cortex and the SCN. UCSC Genome Browser tracks showing differential methylation between Sham and SD for the gene *Gnas* in cortex (shaded blue) and the SCN (shaded pink). The intensity of the filled bar is directly proportional to observed methylation fraction (black = 100% methylation, white = 0% methylation at the CpG site). The chromosome location and scale (mm10 genome) are indicated at the top along with ENCODE tracks for cCRE elements and CpG methylation.

